# NEDD4 promotes Sertoli cell proliferation and adult Leydig cell differentiation in the murine testis

**DOI:** 10.1101/2025.02.19.639149

**Authors:** Simon P. Windley, Yasmine Neirijnck, Diana Vidovic, Quenten Schwarz, Sharad Kumar, Serge Nef, Dagmar Wilhelm

## Abstract

Successful testis development relies on the coordinated differentiation and assembly of various cell types to establish both endocrine and reproductive functions. The ubiquitin ligase NEDD4 has emerged as a key player in murine testis development, with this enzyme being implicated in gonadal sex determination and spermatogonial stem cell differentiation. Here, we report hitherto uncharacterized roles of NEDD4 in postnatal testis development. Utilizing *Nr5a1-* and *Amh-*Cre drivers to conditionally ablate *Nedd4* in testicular somatic cells, we show that NEDD4 promotes Sertoli cell proliferation through the modulation of the PI3K-AKT signaling pathway. This ubiquitin ligase also ensures proper differentiation of adult Leydig cells and may contribute to murine steroidogenesis. Furthermore, NEDD4 is essential for adrenal gland differentiation, as its loss results in adrenal dysgenesis. These findings highlight NEDD4 as a crucial factor in testis development, emphasizing the importance of ubiquitination and post-translational modifications in reproductive biology.

## Introduction

Successful testis development requires the coordinated differentiation and assembly of multiple cell types to enable the endocrine and reproductive capabilities of the mature testis. In mammals, testis fate is determined by the expression of the sex determining region on the Y chromosome (*Sry*) in somatic cell progenitors [1–3]. This initiates a rapid cascade in which SRY up-regulates the expression of a related family member, SRY-box containing gene 9 (*Sox9*), driving the differentiation of the supporting cell lineage into Sertoli cells [4–7]. The critical role of SRY and SOX9 in sex determination is evidenced by complete XY male-to-female gonadal sex reversal upon loss of either factor [8–10] while their over-expression in XX gonads induces testis formation [11, 12].

Once testicular fate is determined, Sertoli cells play a crucial role in orchestrating the differentiation of other testicular cell types, including peritubular myoid cells and steroidogenic fetal Leydig cells [13–17]. They also drive early testicular morphogenesis including testis cord formation and testis vascularization [18–21]. In mice, fetal Leydig cells rapidly increase in number between 12.5 and 15.5 days *post coitum* (dpc) as a result of paracrine recruitment and differentiation of progenitor Leydig cells [16, 22–25]. Fetal Leydig cells ultimately regress postnatally and are replaced by adult Leydig cells. While the origins of fetal and adult Leydig cells are incompletely understood, both Leydig cell types are derived from progenitors within the testis interstitium [26–30]. Adult Leydig cells arise from Leydig stem cells which express *Nestin, Pdgfrb* and *Nr2f2,* which then differentiate into adult Leydig cell progenitors, generally characterized by the upregulation of *Lhr* and expression of *Akr1c14* and *Srd5a1*. Within a week these progenitors proliferate and differentiate into immature Leydig cells, expressing *Hsd17b3*, which increases upon further differentiation into adult Leydig cells [31–34].

Beyond their essential role in fetal development, Sertoli cells remain of crucial importance in the adult testis, where they support and nourish germ cells throughout spermatogenesis [35]. Since each Sertoli cells can sustain only a limited number of germ cells, testis size and sperm production are directly correlated with Sertoli cell number [36]. This Sertoli cell population is established during the late fetal and early perinatal period when they undergo a phase of rapid proliferation, before becoming mitotically inactive at approximately postnatal day (P) 15 [37–40].

The Neural precursor cell Expressed Developmentally Down-regulated protein 4 (NEDD4) is the founding member of the Nedd4 family of ubiquitin protein ligases [41]. Ubiquitin ligases covalently attach ubiquitin or ubiquitin chains to specific substrates. Ubiquitin modifications play essential roles in virtually all aspects of cell signaling through proteolytic and nonproteolytic mechanisms, including proteasomal degradation and protein trafficking. NEDD4 regulates a complex array of pathways during development [42–47] and ablation of NEDD4 in mice results in embryonic lethality [41].

A role for NEDD4 in murine testis development is becoming increasingly evident, as it has been implicated in both gonadal sex determination [48] and spermatogonial stem cell differentiation [49]. To further elucidate the cell autonomous roles of NEDD4 in murine testis development, the Cre/Lox system was employed, allowing for conditional ablation of this E3 ubiquitin ligase specifically in somatic cells of the developing testis. This strategy circumvented the embryonic lethality and growth restriction observed in conventional *Nedd4* constitutive knockout mice [42] enabling spatially restricted ablation of NEDD4 within in the somatic cells of the developing testis.

Here we report that conditional ablation of *Nedd4* in testicular somatic cells did not replicate the sex reversal observed upon constitutive ablation of *Nedd4* [48]. However, it uncovered a later role for NEDD4 in testis development, characterized by a significant reduction in testis size due to impaired Sertoli cell proliferation and disrupted adult Leydig cell differentiation. These results, together with previous studies [48–50], highlight distinct and stage-specific roles of the ubiquitin protein ligase NEDD4 in murine testis development, emphasizing its importance beyond early sex determination.

## Materials and Methods

### Mouse Lines

Conditional *Nedd4* knockout mice, wherein *Nedd4* is ablated in the somatic cells of the developing testis, were generated using *Nedd4^flox/flox^*mice (Nedd4^tm3.1Bros^), in which exon 9 of *Nedd4* is flanked by two *loxP* sites [51], crossed with *Nr5a1*-Cre positive mice (Tg(Nr5a1-cre)2Klp), which drives Cre expression under the *Nr5a1* promoter [52], a gene expressed in the earliest stages of genital ridge formation [53]. *Nr5a1-*Cre*;Nedd4^Flox/+^*mice were time-mated with *Nedd4^Flox/Flox^* mice in order to generate *Nr5a1-Cre;Nedd4^Flox/Flox^* mice, with noon of the day on which a vaginal plug was observed deemed as 0.5 days *post coitum* (dpc). For more accurate staging of fetuses up to 12.5 dpc, the tail somite stage (ts) was determined by counting the number of somites posterior to the hind limb, with 11.5 dpc corresponding with 18 ts, 12.0 dpc corresponding to 24 ts, and 12.5 dpc corresponding to 30 ts [54]. For tissue collected postnatally, noon of the day of birth was determined to be postnatal day (P) 0.

Mice were genotyped using PCR on genomic DNA derived from tail tissue. Primers Nedd4Floxed_F: 5’-*GTACATTTTAGTTCATGGTTCTCACAGG*-3’ and Nedd4Floxed_R: 5’-*CAGAGCTCACATGGCTGTGGG*-3’ resulted in PCR products of 168 base pairs (bp) and 202 bp for the wildtype (WT) and floxed alleles respectively, while primers Cre408_F: 5’-*GCATTACCGGTCGATGCAACGAGTGATGAG*-3’ and Cre408_R: 5’-*GAGTGAACGAACCTGGTCGAAATCAGTGCG*-3’ generated a 408 bp product in Cre-positive mice. Genetic sex was determined as described previously [55].

For Sertoli cell specific ablation of *Nedd4, Nedd4^flox/flox^* mice (Nedd4^tm3.1Bros^) were mated with mice containing the *Amh*-Cre transgene (Tg(AMH-cre)1Flor) [56] to generate *Amh-*Cre*;Nedd4^Flox/Flox^* mice, which were genotyped using the Nedd4Floxed primers described above and Cre26: 5’-*CCTGGAAAATGCTTCTGTCCG*-3’ and Cre36: 5’-*CAGGGTGTTATAAGCAATCCC*-3’, which amplified a 400 bp Cre fragment.

All animal experiments were performed with the approval of the Animal Ethics Committee of the University of Melbourne (approval number 1513725).

### Immunofluorescence

Section immunofluorescence on paraformaldehyde (PFA)-fixed, paraffin embedded mouse fetuses was performed as described previously [50]. Primary antibodies used were mouse anti-NEDD4 (BD Transduction Laboratories, 611481; 1:100), goat anti-DDX4 (R&D Systems, RDSAF2030; 1:300), rabbit anti-SOX9 [6]1:200), rabbit anti-SRY [6] 1:100), rabbit anti-FOXL2 [57](1:300), goat anti-GATA4 (Santa Cruz, sc1237; 1:300), rabbit anti-Laminin (Sigma Aldrich, L9393; 1:300), rabbit anti-CYP11A1 [58]1:300), mouse anti-POU5F1 (Santa Cruz, sc5279; 1:100), mouse anti-SYCP3 (Abcam, ab97672; 1:100), goat anti-AMH (Santa Cruz, sc6886; 1:300), sheep anti-SOX9 [59] 1:200), rat anti-NR5A1 (Transgenic Inc, KO610; 1:500), rabbit anti-TH (Sigma-Aldrich AB152; 1:100), mouse anti-Ki67 (BD Transduction Lab, 550609; 1:100) and mouse anti-NR2F2 (Perseus Proteomics, PP-H7147-00; 1:200). All secondary antibodies were purchased from Invitrogen and were used at a dilution of 1:300.

### Quantitative real-time RT-PCR (RT-qPCR)

Mouse fetuses were collected at 14.5 dpc, gonad pairs were dissected and the underlying mesonephros removed before being snap-frozen in liquid nitrogen. Similarly, whole testes were dissected from P60 mice and snap-frozen in liquid nitrogen. Total RNA was isolated using Trizol (Ambion) and single stranded cDNA synthesized using the Protoscript II First Strand cDNA Synthesis Kit (New England Biolabs). qPCR was performed with the SensiFAST SYBR No-ROX Kit (Bioline) using the Rotor-Gene 3000 system (Qiagen). Relative mRNA levels were normalized to *Sdha* (fetal) or *Tbp* (postnatal) and are shown relative to controls. Each reaction was performed in technical triplicate and each experiment was performed with at least five biological replicates. N numbers are specified in each corresponding figure legend. Values are plotted as the mean, with error bars representing the standard error of the mean (SEM) between biological replicates. Statistical significance between groups was determined using a two-tailed, unpaired Student’s t-test. Primer sequences can be found in Table S1.

### Testis and seminal vesicle weight measurement

Testes of P7, P14, P28 and P60 (*Nr5a1-*Cre*;Nedd4^Flox/Flox^*) and P15 and P60 (*Amh-* Cre*;Nedd4^Flox/Flox^*) and seminal vesicles of P60 *Nr5a1-Cre;Nedd4^Flox/Flox^*and *Amh-* Cre*;Nedd4^Flox/Flox^* control and mutant animals were dissected from each mouse and weighed. Weights were normalized to body weight and are displayed as the mean ± SEM with the average of controls set to 100%. Statistical significance between groups was determined using a two-tailed, unpaired Student’s t-test.

### Histology

Testes were fixed overnight in Bouin’s fixative and embedded in paraffin. To evaluate the histology of these organs, 5 μm transverse sections were stained with hematoxylin and eosin (H&E).

### Sertoli cell proliferation assay

Paraformaldehyde-fixed, paraffin embedded 15.5 dpc *Nr5a1-*Cre*;Nedd4^Flox/Flox^*(n = 5) and control littermate fetuses (n = 4) were sectioned and co-immunolabelled with rabbit anti-SOX9 and mouse anti-Ki67. Between 3-5 whole testis cross sections, at least 50 μm distant from one another were selected and imaged on a Zeiss, LSM800 confocal microscope. Sections were manually counted by blinded observers.

Proliferating Sertoli cells (KI67^+^/SOX9^+^) are shown as a percentage of total Sertoli cells (SOX9^+^). Statistical significance between groups was determined using a two-tailed, unpaired Student’s t-test.

### Immunoblotting

P60 mouse testes were homogenized, lysed in RIPA buffer (50 mM Tris-HCL [pH 7.5], 150 mM NaCl, 1% NP-40, 1mM EDTA, 1 mM EGTA, 0.1% SDS and 0.5% Sodium deoxycholate) supplemented with cOmplete^TM^ Protease Inhibitor Cocktail (Roche) and total protein measured using Bradford Assay (Thermo Scientific) before being further diluted in RIPA buffer as required. 4x SDS loading buffer (200mM Tris-HCl [pH 6.8], 0.4M DTT, 8% SDS, 0.2% bromophenol blue, 40% glycerol) was added to each sample and boiled for 10 minutes before separating on 7.5-, 10-or 12% polyacrylamide gels for 2 hours at 100V. Separated protein samples were transferred onto polyvinylidene difluoride membranes, washed 3 times with Tris buffered saline supplemented with 0.1% Tween^®^ 20 detergent (TBST) and blocked with 5% bovine serum albumin in TBST for 1 hour. Membranes were incubated with primary antibody diluted 1:1000 in blocking solution overnight at 4°C, washed 3 times in TBST and incubated with a HRP coupled secondary antibody for two hours. Membranes were washed 3 more times with TBST, prepared for imaging through the addition of a 1:1 mixture of H_2_O_2_ and Luminol (SuperSignal West Pico PLUS Kit, Thermo Fisher Scientific) and imaged using a ChemiDoc MP (BioRad). Primary antibodies used in this study are mouse anti-NEDD4 (BD Transduction Laboratories, 611481), rabbit anti-PTEN (Cell Signaling Technology, 9559), rabbit anti-phospho-AKT (Ser473) (Cell Signaling Technology, 9271), rabbit anti-β-Actin (Cell Signaling Technology, 4967), rabbit anti-Beclin (Cell Signaling Technology, 3495S), rabbit anti-p62 (Cell Signaling Technology, 39749), rabbit anti-LC3B (Cell Signaling Technology, 3868), rabbit anti-p53 (Cell Signaling Technology, 32532), mouse anti-PDCD6IP (aka. ALIX) (Cell Signaling Technology, 92880) and rabbit anti-GAPDH (Cell Signaling Technology, 21185).

### Sperm count and sperm motility

For *Amh-*Cre*;Nedd4^Flox/Flox^* mice cauda epididymides and vas deferens of P60 males were harvested in prewarmed M2 media, and fat and connective tissues were removed. Sperm cells were released in 1mL of M2 media by manual dissociation of the tissue followed by an incubation of 10 min at 37°C. A 1:40 dilution of the sperm suspension was loaded in a prewarmed 100µm-deep counting chamber (Leja Products, Nieuw-Vennep, The Netherlands) and submitted to computer-assisted sperm analysis (CASA) (CEROS II, Hamilton Thorn Research Inc, Beverly MA). The settings employed for analysis were as follows: acquisition rate: 60 Hz; number of frames: 100; minimum illumination: 70; maximum illumination: 90; minimum cell area: 8; maximum cell area: 100; minimum elongation gate: 5; maximum elongation gate: 90; minimum cell brightness: 129; minimum tail brightness: 102; magnification factor: 0.74. The motility parameters measured were curvilinear velocity (VCL), average path velocity (VAP) and amplitude of lateral head displacement (ALH). Static, motile, progressive and hyperactivated sperm were characterized by the default parameters of the software. At least 300 sperm cells were analyzed for each assay. The number of sperm present in the initial 1 mL suspension was calculated by the CASA system (10^6^ sperm cells/mL).

## Results

### Fetal testis development proceeds normally in Nr5a1-Cre;Nedd4^Flox/Flox^ mice

Previous studies identified a role for NEDD4 in murine gonadal sex determination, where constitutive *Nedd4* ablation led to complete male-to-female sex reversal in XY mice [48]. To further investigate the cell autonomous roles of NEDD4 within the somatic gonad, while circumventing the fetal lethality associated with constitutive ablation, we employed the Cre-Lox system to conditionally delete *Nedd4* in gonadal somatic cells, utilizing Cre recombinase under the control of the *Nr5a1* promoter (*Nr5a1-*Cre*;Nedd4^Flox/Flox^* mice).

In *Nedd4*-null gonads, sex reversal is characterized by insufficient progenitor cell proliferation, delayed onset of SRY expression, negligible SOX9 levels and ectopic activation of the ovarian program [48]. In contrast, immunofluorescence analysis of *Nr5a1-*Cre*;Nedd4^Flox/Flox^* and control gonads at E11.5 revealed normal SRY and SOX9 expression in Sertoli cells, along with DDX4-positive germ cells (Figure 1A). Although a few FOXL2-positive cells were observed in *Nr5a1-*Cre*;Nedd4^Flox/Flox^*gonads, they were insufficient to promote ovarian fate. By 12.5 dpc, SRY expression was down-regulated and SOX9-positive Sertoli cells had upregulated anti-Müllerian hormone (AMH) and formed testis cords with germ cells (Figure 1B). Despite an apparently normal testis-determining program, *Nr5a1-*Cre*;Nedd4^Flox/Flox^*testes were slightly smaller than control by 12.5 dpc (Figure 1B, bottom panel). Taken together, these findings indicated that XY *Nr5a1-*Cre*;Nedd4^Flox/Flox^* mice did not replicate the sex reversal observed upon constitutive ablation of *Nedd4*.

**Figure 1.**
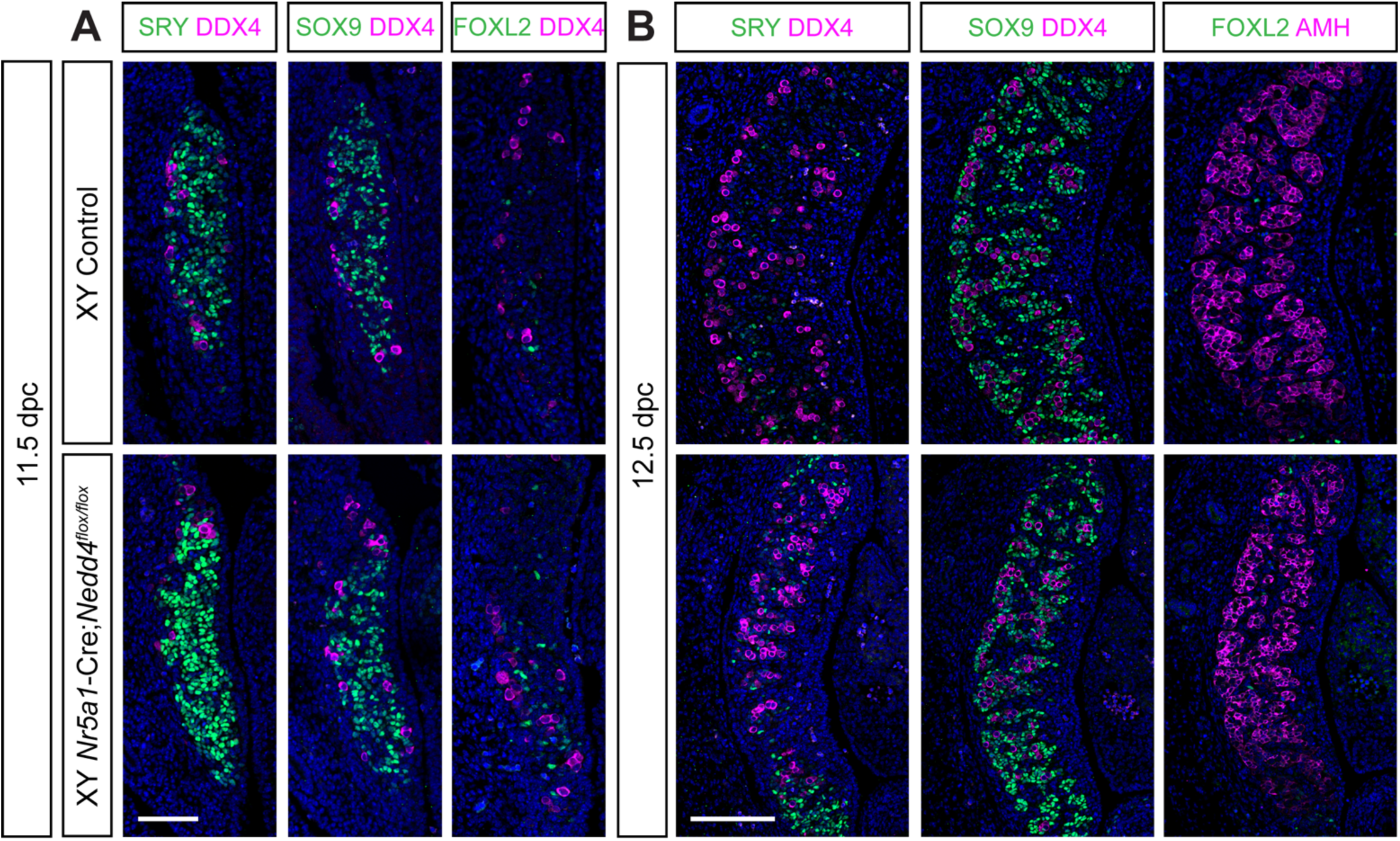
The Testis Determining Program is Unaffected in XY *Nr5a1-*Cre*;Nedd4^Flox/Flox^* Mice. Section immunofluorescence on 11.5 dpc (**A**) and 12.5 dpc (**B**) XY *Nr5a1*-Cre*;Nedd4^Flox/Flox^* gonads (bottom) alongside Cre-negative XY littermate controls (top) stained for Sertoli cell markers SRY (green, left panel), SOX9 (green, middle panel) and AMH (magenta right panel in **B**), granulosa cell marker FOXL2 (green, right panel) and germ cell marker DDX4 (magenta in A and left and middle panels of **B**). The anterior pole of each gonad is positioned at the top of each panel. Scale bars = 100 μm.

To investigate the discrepancy in gonadal phenotypes between *Nr5a1-* Cre*;Nedd4^Flox/Flox^* and *Nedd4-*null mice, and to confirm *Nedd4* ablation, we performed NEDD4 immunofluorescence alongside the germ cell marker DDX4 at 11.5 dpc, approximately 32 hours after NR5A1 is first expressed [60]. While NEDD4 was ubiquitously expressed throughout the gonad and surrounding tissues of control littermates, *Nr5a1-*Cre*;Nedd4^Flox/Flox^* mice exhibited a reduction in NEDD4 expression in most DDX4-negative somatic gonadal cells, although expression was still observed in some somatic cells within the gonad (Figure S1A). Some of these cells may be endothelial cells, as these are not derived from NR5A1-positive progenitors. Unexpectedly, NEDD4 expression remained detectable in coelomic epithelial cells of XY *Nr5a1-*Cre*;Nedd4^Flox/Flox^*mice, indicating that *Nedd4* ablation was incomplete by 11.5 dpc. At 14.5 dpc, *Nedd4* transcripts were significantly reduced (Figure S1B), yet NEDD4 protein was still present within XY *Nr5a1-Cre;Nedd4^Flox/Flox^* gonads, albeit at a much lower level that in controls. While NEDD4 was widely expressed in control gonads (Figure S1C), consistent with our previous studies [48, 50], it was largely absent in SOX9-positive Sertoli cells and interstitial Leydig cells in XY *Nr5a1-* Cre*;Nedd4^Flox/Flox^* mice (Figure S1C). While NEDD4 expression in germ cells within testis cords of *Nr5a1-*Cre*;Nedd4^Flox/Flox^*mice was expected (Figure S1C, white box), due to their *Nr5a1-*independent origins, the persistence of NEDD4 within the coelomic epithelium was unexpected (Figure S1C, arrow heads). Given that sex reversal in XY *Nedd4^-/-^* gonads is attributed to insufficient proliferation of the coelomic epithelium, leading to a reduced pre-Sertoli cell pool, it is plausible that residual NEDD4 expression within the coelomic epithelium in XY *Nr5a1-* Cre*;Nedd4^Flox/Flox^*gonads is sufficient to overcome this proliferation deficit, allowing the testis-determining program to proceed unperturbed.

Although XY *Nr5a1-*Cre*;Nedd4^Flox/Flox^* mice did not replicate the gonadal sex reversal seen in XY *Nedd4^-/-^* mice, they provided a valuable model to investigate the role of NEDD4 in the somatic cells of the developing testis, a task previously hindered by the complete gonadal sex reversal of XY *Nedd4*^-/-^ mice. To explore this, we performed immunofluorescence on 14.5 dpc XY *Nr5a1-* Cre*;Nedd4^Flox/Flox^* gonads, assessing key testicular markers (SOX9, GATA4, CYP11A1, AMH) and ovarian marker (FOXL2) to determine somatic cell identity. Additionally, we examined germ cell markers (DDX4, POU5F1, SYCP3) to evaluate potential indirect effects of somatic cells in instructing the development of fetal germ cells. Our analysis revealed that the testes of XY *Nr5a1-*Cre*;Nedd4^Flox/Flox^* mice were morphologically and molecularly similar to Cre-negative littermate controls. Sertoli cells expressing SOX9 (Figure 2A, F), GATA4 (Figure 2B, G) and AMH (Figure 2D, I), successfully formed testis cords which enclosed DDX4-positive germ cells (Figure 2A, F, E, J) and were outlined by the extracellular matrix protein laminin (Figure 2B, G). Steroidogenic fetal Leydig cells, marked by CYP11A1 (Figure 2C, H), properly differentiated within in the testis interstitium. No FOXL2-positive cells, indicative of gonadal sex reversal, were observed (Figure 2E, J). Finally, while gonadal sex reversal extended to the developing germ cells in *Nedd4^-/-^* mice [48], germ cells differentiated in line with that expected of gonocytes within the developing testis, retaining expression of the pluripotency marker POU5F1 (Figure 2C, H) and staining negative for SYCP3 (Figure 2D, I), a meiosis marker typical of germ cells within the developing ovary at this age.

**Figure 2.**
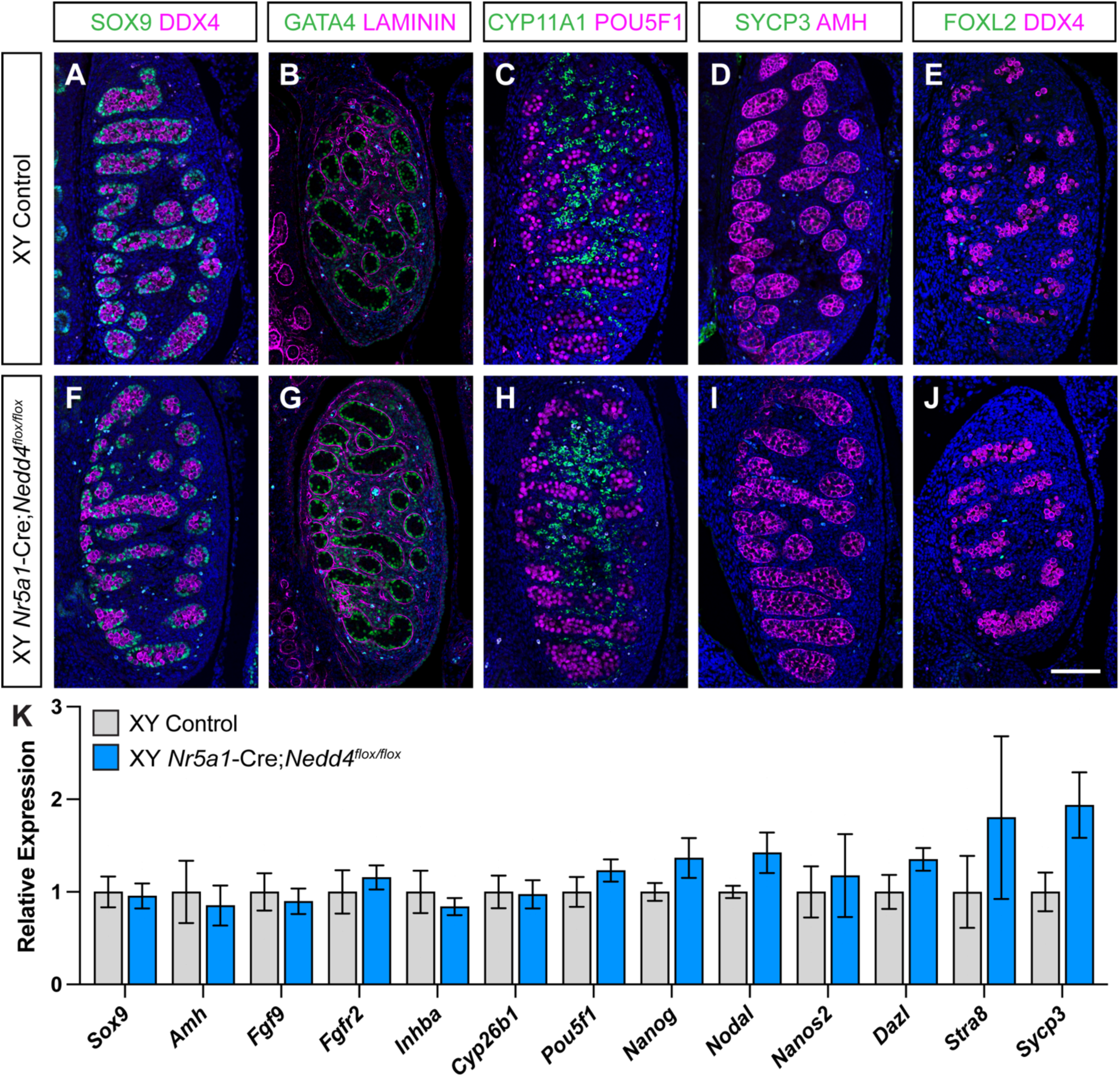
Unperturbed Fetal Testis Differentiation in XY *Nr5a1-*Cre*;Nedd4^Flox/Flox^* Mice. Section immunofluorescence on 14.5 dpc XY *Nr5a1-*Cre;*Nedd4^flox/flox^*testes alongside XY littermate controls stained for Sertoli cell markers SOX9 (green in **A**, **E**), GATA4 (green in **B**, **G**) and AMH (magenta in **D**, **I**), extracellular matrix protein laminin (magenta in **B**, **G**), Leydig cell marker CYP11A1 (green in **C**, **H**), germ cell marker DDX4 (magenta in **A**, **E**, **F**, **J**), pluripotency marker POU5F1 (magenta in **C**, **H**), meiosis marker SYCP3 (green in **D**, **I**) and granulosa cell marker FOXL2 (green in **E**, **J**). The anterior pole of each gonad is positioned at the top of each panel. Scale bars = 100 μm. **K**) RT-qPCR analyses of *Sox9, Amh, Fgf9, Fgfr2, Inhba, Cyb26b1, Pou5f1, Nanog, Nodal, Nanos2, Dazl, Stra8* and *Sycp3* expression at 14.5 dpc on XY *Nr5a1-* Cre;*Nedd4^flox/flox^*gonads (blue, n=5) and XY littermate controls (grey, n=5). Values are normalized to *Sdha* and are expressed relative to controls. Mean ± SEM; t-test; n.s. = not significant.

RT-qPCR analysis of micro-dissected tissue further confirmed that testis development remained unperturbed in *Nr5a1-*Cre*;Nedd4^Flox/Flox^*testes (Figure 2K). Mutant testes showed no significant differences in transcripts associated with Sertoli cells (*Sox9, Amh, Fgf9, Fgfr2, Inhba* and *Cyp26b1*), male embryonic germ cell development (*Pou5f1, Nanog, Nodal* and *Nanos2*) or female germ cell markers indicative of gonadal sex reversal (*Dazl, Stra8* and *Sycp3*) (Figure 2K). These findings confirm that by 14.5 dpc, *Nr5a1-*Cre*;Nedd4^Flox/Flox^* develop normally, without deviation from control.

Taken together, these data suggest that, beyond its crucial role in establishing gonadal precursors, NEDD4 is not required for the maintenance of Sertoli or fetal Leydig cell identity.

### Adrenal cortex dysgenesis in Nr5a1^Cre/+^;Nedd4^Flox/Flox^ mice

Given the shared developmental origins of the gonads and adrenal glands [53, 61], and the expression of *Nr5a1-*Cre in both gonadal somatic cells and the adrenal cortex [52], we also examined adrenal gland development in *Nr5a1-*Cre*;Nedd4^Flox/Flox^*mice. In Cre-negative animals, NEDD4 was strongly expressed in the cytoplasm of NR5A1-positive cells of the adrenal cortex but was largely absent from the neural-crest derived chromaffin cells of the adrenal medulla at 14.5 dpc. In contrast, while NEDD4 expression was still observed in surrounding tissues of *Nr5a1-* Cre*;Nedd4^Flox/Flox^* mice, expression was absent from the NR5A1-positive cell population, confirming successful ablation of *Nedd4* in the adrenal cortex (Figure S2A). Notably, in both XX and XY *Nedd4-*ablated adrenal glands, the cortical cell population was significantly reduced compared to Cre-negative controls (Figure S2), suggesting an essential role for NEDD4 in adrenal cortex development.

To visualise this phenotype further, immunofluorescence staining was performed on *Nr5a1-*Cre*;Nedd4^Flox/Flox^* and control embryos at 14.5, 16.5 and 18.5 dpc using antibodies for NR5A1 and the chromaffin cell marker tyrosine hydroxylase (TH) [62]. This analysis confirmed that *Nedd4-*deficient adrenal glands exhibited a severely diminished NR5A1-positive cortical cell population, resulting in a hypoplastic adrenal cortex and a failure to fully enclose the medulla (Figure S2B). From 14.5 dpc the developing adrenal gland becomes encapsulated by fibrous mesenchyme-derived tissue, establishing distinct cortical and medullary compartments, a process that is nearly completed by birth [63, 64]. In Cre-negative controls, a well-defined adrenal cortex progressively expanded around the medulla, with these cell populations appearing near mutually exclusive by 18.5 dpc. In contrast, while NR5A1-positive adreno-cortical cells in *Nr5a1-*Cre*;Nedd4^Flox/Flox^*embryos increased over time, their numbers remained insufficient to fully encapsulate the chromaffin cell population, resulting in a disorganized cortex-medulla boundary (Figure S2B). Collectively, these data show that NEDD4 is expressed within the NR5A1+ cell population of the adrenal cortex and is required for adrenal cortex development, with the absence of *Nedd4* resulting in adrenal cortex dysgenesis.

### Aberrant Sertoli cell proliferation upon loss of Nedd4

To further investigate the role of NEDD4 in testicular somatic cell development, we analyzed adult *Nr5a1-*Cre*;Nedd4^Flox/Flox^*testes at postnatal day 60 (P60). Mutant testes exhibited a significant reduction in size (Figure 3A, right testis) with testis weight decreased by 47.8% compared to littermate controls (Figure 3B). Despite this size reduction, histological analysis using haematoxylin and eosin staining revealed little difference in the seminiferous epithelium of *Nr5a1-*Cre*;Nedd4^Flox/Flox^* testes when compared with controls, with both containing seminiferous tubules capable of supporting spermatogenesis, as indicated by the presence of spermatozoa within the tubule lumen (Figure 3C, D). Immunofluorescence analysis for SOX9 and FOXL2 at P60 further confirmed that Sertoli cells maintained their identity, with SOX9-positive Sertoli cells lining the basement membrane of seminiferous tubules, while the ovarian program was repressed, as evidenced by the absence of FOXL2 expression (Figure 3E, F). To determine when testis size differences first emerged, we assessed testis weight at P7, P14 and P28. At all ages *Nr5a1-*Cre*;Nedd4^Flox/Flox^*testes were significantly smaller than controls, exhibiting reductions of 39.5% at P7, 48.6% at P14 and 38.7% at P28 (Figure 3G).

**Figure 3.**
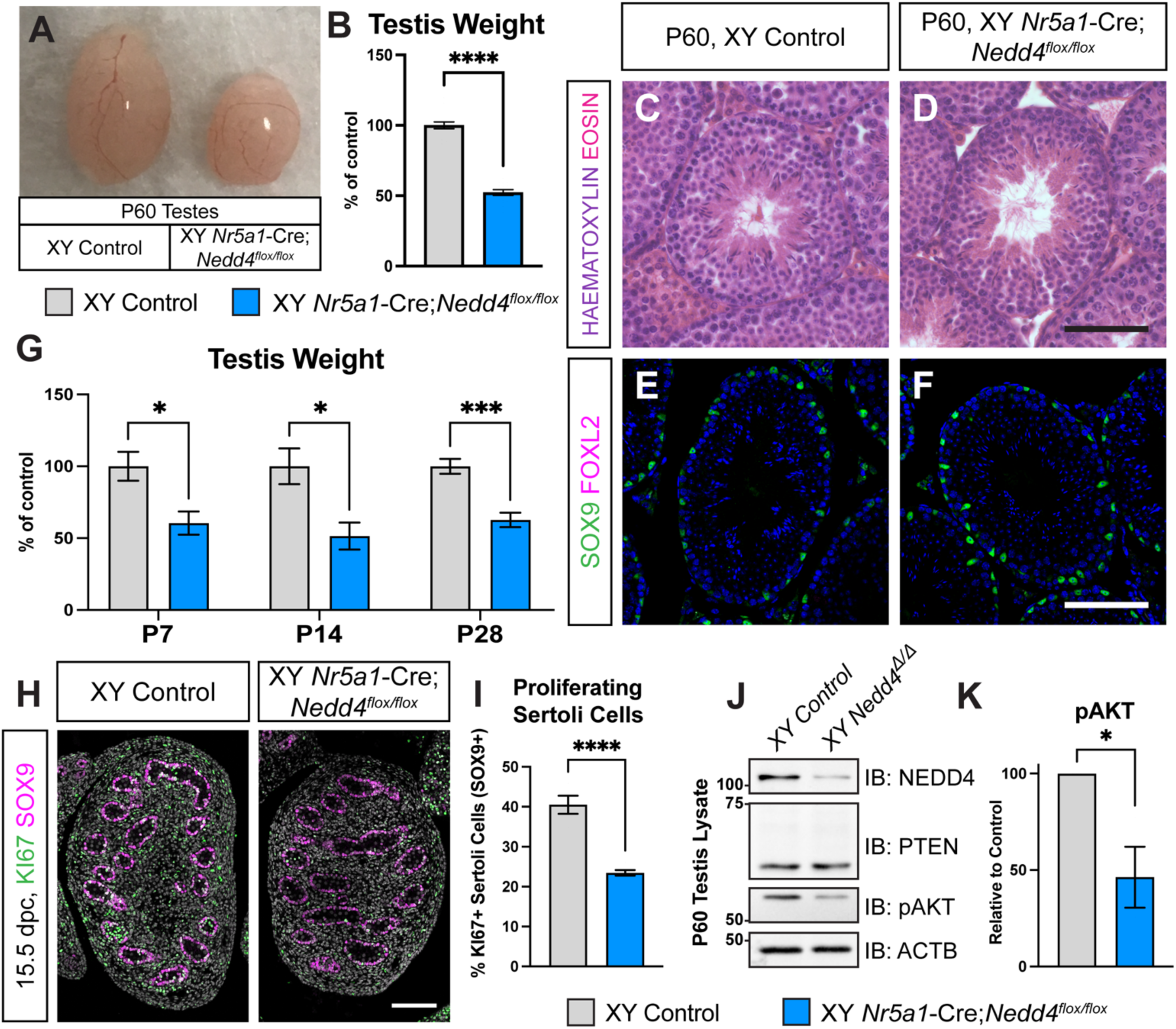
Impaired Sertoli Cell Proliferation in *Nr5a1-*Cre;*Nedd4^flox/flox^* Mice. Testes **(A)** and Testis Weight (**B**, n=14) of postnatal day (P) 60 Cre-negative littermate control (left, grey) and *Nr5a1*-Cre;*Nedd4^flox/flox^* (right, blue) mice. Haematoxylin and Eosin staining of Cre-negative (**C**) and *Nr5a1-*Cre;*Nedd4^flox/flox^* (**D**) P60 testes. Section immunofluorescence on P60 Cre-negative control (**E**) and *Nr5a1-*Cre;*Nedd4^flox/flox^*(**F**) testes stained for Sertoli cell marker SOX9 (green), granulosa cell marker FOXL2 (magenta). Scale bars = 100 μm. **G**) Testis weights of control (grey) and *Nr5a1-*Cre;*Nedd4^flox/flox^* mice (blue) at P7 (n = 5), P14 (n= 6 control, 5 mutant) and P28 (n = 11 control, 9 mutant). Testis weights are normalized to body weight and are shown as a percentage of controls. **H**) Section immunofluorescence on 15.5 dpc control and *Nr5a1-*Cre;*Nedd4^flox/flox^* testes stained for proliferation marker KI67 (green) and Sertoli cell marker SOX9 (magenta), counterstained with DAPI (grey). Scale bars = 100 μm. **I**) Quantification of Sertoli cell proliferation of control (grey, n = 4) and *Nr5a1-*Cre;*Nedd4^flox/flox^* (blue, n = 5) fetal testes, shown as the proportion of proliferating Sertoli cells (KI67+ and SOX9+) within the total Sertoli cell pool (SOX9+). **J**) Immunoblot on control (left) and *Nr5a1-* Cre;*Nedd4^flox/flox^* (*Nedd4^Δ/Δ^*, right) P60 testis lysates probed with NEDD4, PTEN, phosphorylated AKT (pAKT) and Beta Actin (ACTB). Size markers (kilo Daltons) are shown on the lefthand side. **K**) Quantification of pAKT immunoblot of control (grey, n = 3) and *Nr5a1-*Cre;*Nedd4^flox/flox^* (blue, n = 3) P60 testis lysates. Values are normalized to Beta Actin and are shown relative to littermate testes. All graphs display mean ± SEM; t-test; * = p<0.05, *** = p<0.001, **** = p<0.0001.

Since testis size is directly correlated with the total number of adult Sertoli cells, (reviewed in Sharpe, McKinnell [65]) and reduced Sertoli cell proliferation leads to decreased testis weight [37], we next assessed the proliferative capacity of fetal Sertoli cells, to determine whether diminished Sertoli cell proliferation could contribute to the postnatal weight reduction observed in *Nr5a1-*Cre*;Nedd4^Flox/Flox^*testes. Indeed, at 15.5 dpc, fetal *Nr5a1-*Cre*;Nedd4^Flox/Flox^*testes exhibited 42.1% fewer proliferative Sertoli cells (KI67+ and SOX9+) compared to littermate controls (Figure 3H, I). NEDD4 is recognized for its oncogenic potential through the negative regulation of the tumour suppressor phosphatase and tensin homolog (PTEN), a well characterized negative regulator of the PI3K-Akt signaling pathway [66]. Since NEDD4 and PTEN levels are often inversely correlated [67–69] and PTEN has been shown to negatively regulate Sertoli cell proliferation [70], we assessed whether increased PTEN expression, and diminished PI3K-AKT signaling might explain the reduced Sertoli cell proliferation in *Nedd4-*mutant testes. As expected, NEDD4 protein levels were significantly reduced in P60 *Nr5a1-*Cre*;Nedd4^Flox/Flox^*testes but mutant testes showed no significant differences in total PTEN levels (Figure 3J). Despite this, signaling along the PI3K-AKT axis was still diminished, with a 53.7% reduction of phosphorylated AKT (Figure 3J, K). Together, these findings suggest that NEDD4 promotes Sertoli cells proliferation by regulating PI3K-AKT signaling, and its loss leads to impaired Sertoli cell proliferation and reduced testis size.

To further explore the role of NEDD4 in Sertoli cells, we generated *Amh-*Cre*;Nedd4^Flox/Flox^* mice, in which *Nedd4* was selectively ablated in Sertoli cells from approximately 15 dpc [56]. While *Nr5a1-*Cre*;Nedd4^Flox/Flox^* testes were significantly smaller than those of their littermates as early as P7, this was not the case upon Sertoli cell specific ablation of *Nedd4*, with *Amh-*Cre*;Nedd4^Flox/Flox^*mutant testes being comparable in weight and size to controls at P15. By P60, however, *Amh-* Cre*;Nedd4^Flox/Flox^* testes showed a 29.7% reduction in weight compared to littermates (Figure 4A, B). At P60, mutant testes frequently displayed abnormal morphology, appearing shorter in length and often associated with a larger blood vessel (Figure 4C, red arrows) in contrast to littermates (Figure 4C, white arrows). Similar to *Nr5a1-* Cre*;Nedd4^Flox/Flox^* mice, histological analysis of mutant testes revealed no major alteration of the seminiferous epithelium (Figure 4D). *Amh-*Cre*;Nedd4^Flox/Flox^* mice were viable, reached adulthood and exhibited normal sexual behaviour, internal and external genitalia and unaltered seminal vesicles weights (Figure 4E).

**Figure 4.**
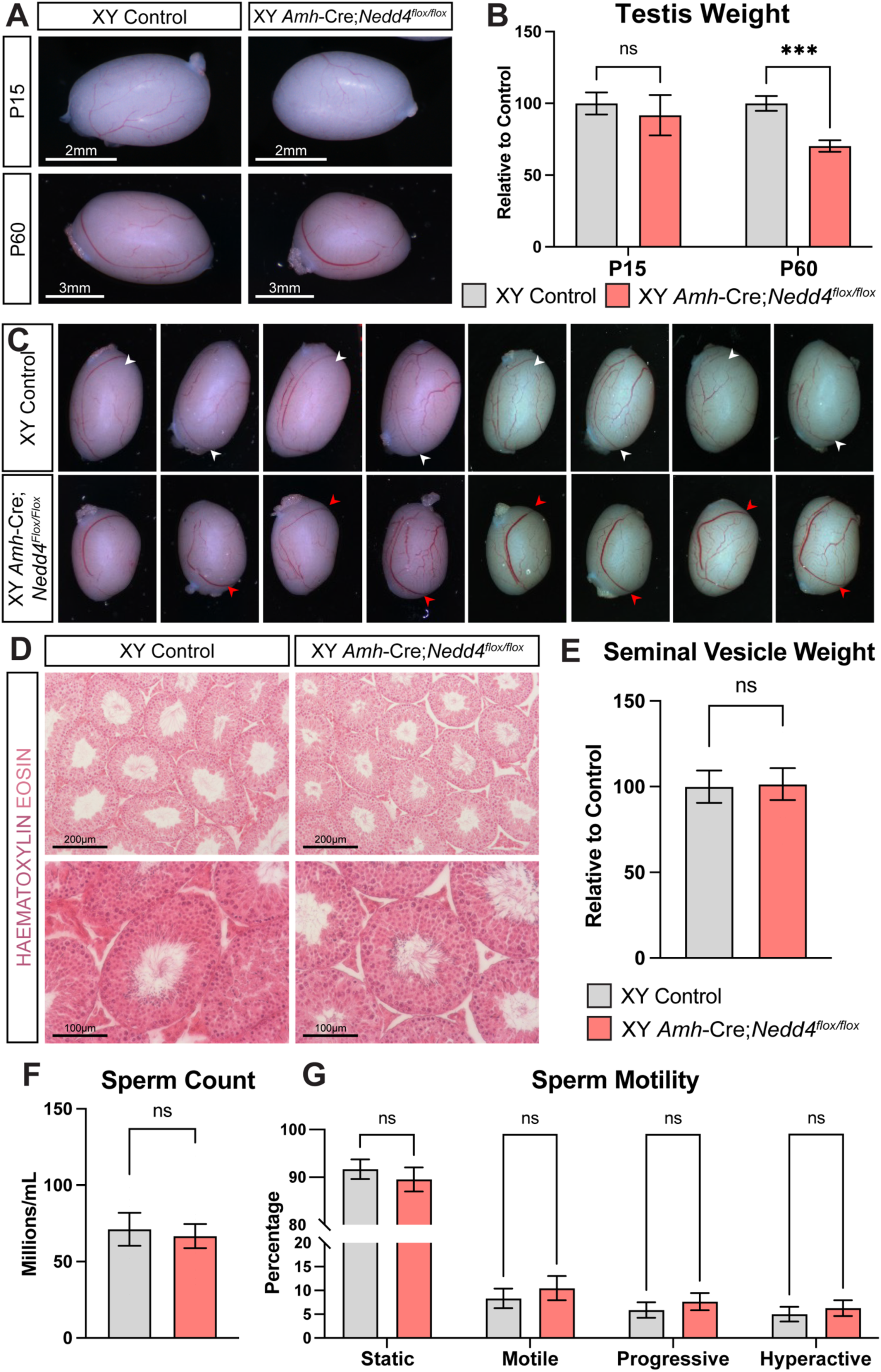
Reduced Testis Size and Abnormal Testis Morphology Upon Sertoli Cell Specific Ablation of *Nedd4*. Testes (**A**) from control (left) and Sertoli cell specific *Nedd4* mutants (*Amh-* Cre;*Nedd4^flox/flox^,* right) at postnatal (P) day 15 (top) and P60 (bottom). **B**) Testis Weight of P15 and P60 Cre-negative littermate controls (left, grey, n = 9 [P15], n = 7 [P60]) and *Amh*-Cre;*Nedd4^flox/flox^* mice (right, red, n = 10 [P15], n = 11 [P60]) normalized to body weight and shown as a percentage of controls. **C**) Testes of P60 controls (upper panel) and *Amh*-Cre;*Nedd4^flox/flox^* mice (lower panel) showing abnormal large blood vessels (red arrowhead) in mutants compared to their control littermates (white arrowhead). **D**) Haematoxylin and Eosin staining of P60 control and *Amh-*Cre;*Nedd4^flox/flox^* testes. Seminal vesicle weight (n = 5 control, 8 mutant) (**E**), sperm count (n = 6 control, 10 mutant) (**F**) and sperm motility measures (n = 6 control, 10 mutant) (**G**) of P60 control (left, grey) and *Amh*-Cre;*Nedd4^flox/flox^* mice (right, red). All graphs display mean ± SEM; t-test; n.s. = not significant, *** = p<0.001.

The reduction in testis weight, upon Sertoli cell specific loss of *Nedd4,* was not associated with impaired sperm production. Computer-assisted-sperm-analysis (CASA) revealed no significant differences in sperm counts between mutants and controls (Figure 4F). Additionally, sperm produced by *Amh-*Cre*;Nedd4^Flox/Flox^* mice exhibited normal levels of motility as those produced by their littermates (Figure 4G). These data suggest that the absence of NEDD4 in Sertoli cells only partially contributes to the phenotype observed in *Nr5a1-*Cre*;Nedd4^Flox/Flox^*mice, where *Nedd4* is deleted in almost all testicular somatic cells.

### Aberrant Leydig cell differentiation contributes to reduced testis size in Nr5a1-Cre;Nedd4^Flox/Flox^ testes

Since *Amh*-Cre;*Nedd4^Flox^*^/*Flox*^ mice did not fully replicate the phenotypes observed in *Nr5a1-*Cre*;Nedd4^Flox/Flox^* mice, we next assessed whether defective Leydig cell development or function, could contribute to the testis size reduction. While fetal Leydig cell differentiation seemingly occured as expected (Figure 2), we hypothesized that impaired adult Leydig cell differentiation might exacerbate the observed reduction in testis weight. immunofluorescence at P60 for CYP11A1, a steroidogenic marker that increases with Leydig cell differentiation [31, 71], and NR2F2, a marker for both fetal [72] and adult [29] Leydig progenitor cells, revealed that while *Nr5a1-*Cre*;Nedd4^Flox/Flox^*testes contained both progenitor and differentiated Leydig cells, their interstitial spaces appeared less cellularly dense compared to controls (Figure 5A, B). This is also apparent in haematoxylin and eosin stained sections (Figure 3C, D). To confirm whether perturbed adult Leydig cell differentiation contributes to this diminished cellularity, we next utilized RT-qPCR to assess the relative expression of transcripts of adult Leydig cells at various states of differentiation.

**Figure 5.**
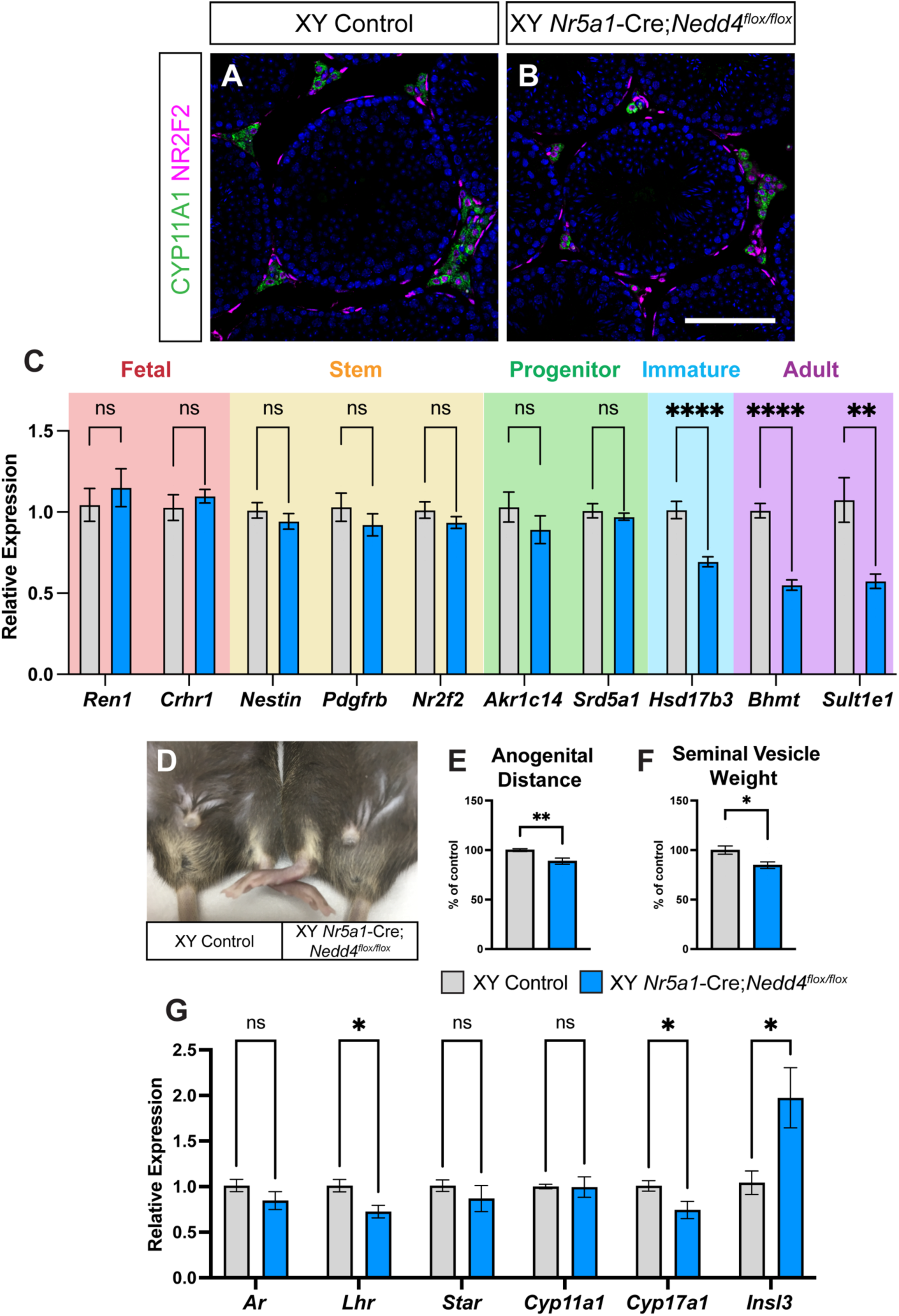
Perturbed Adult Leydig Cell Differentiation in *Nr5a1*-Cre;*Nedd4^flox/flox^* Testes. Section immunofluorescence on P60 Cre-negative control (**A**) and *Nr5a1*-Cre;*Nedd4^flox/flox^* (**B**) testes stained for Leydig cell markers CYP11A1 (green) and NR2F2 (magenta). Scale bars = 100 μm. **C**) RT-qPCR analyses for fetal Leydig cell transcripts *Ren1* and *Crhr1* (red section), Leydig stem cell transcripts *Nestin, Pdgfrb* and *Nr2f2* (yellow section), Leydig progenitor cell transcripts *Akr1c14* and *Srd5a1* (green section), immature Adult Leydig cell transcript *Hsd17b3* (light blue section) and Adult Leydig cell transcripts *Bhmt* and *Sult1e1* (purple section) on P60 control (grey, n= 10) and *Nr5a1-*Cre;*Nedd4^flox/flox^* testes (blue, n=9). Anogenital Distance (**D**, **E**, n = 7) and Seminal Vesicle Weight (**F**, n = 11 control, 10 mutant) of P60 Cre-negative littermate control (Left, grey) and *Nr5a1*-Cre;*Nedd4^flox/flox^* mice (Right, blue). Seminal vesicle weight and anogenital distance are normalized to body weight and are shown as a percentage of controls. **F**) RT-qPCR analyses for expression of genes encoding *Ar, Lhr, Star, Cyp11a1, Cyp17a1* and *Insl3* at P60 in *Nr5a1-*Cre;*Nedd4^flox/flox^* testes (blue, n=5) and Cre-negative littermate controls (grey, n=6). Values are normalized to *Tbp* and are expressed relative to controls. Mean ± SEM; t-test; n.s. = not significant, * = p<0.05, ** = p<0.01, **** = p<0.0001.

This analysis revealed that, while transcripts indicative of fetal Leydig cells (*Ren1* and *Crhr1*), Leydig stem cells (*Nestin, Pdgfrb* and *Nr2f2*) and progenitor adult Leydig cells (*Akr1c14* and *Srd5a1*) were comparable between *Nr5a1-*Cre*;Nedd4^Flox/Flox^*testes and controls at P60, transcripts representing immature and adult Leydig cells (*Hsd17b3, Bhmt* and *Sult1e1*) were significantly reduced in the *Nedd4-*mutant testes (Figure 5C), in line with a seeming reduction in the adult Leydig cell population. Consistent with defective steroidogenesis, *Nr5a1-*Cre*;Nedd4^Flox/Flox^* mice displayed reduced anogenital distance (Figure 5D, E, 11.2% reduction) and decreased seminal vesicle weight (Figure 5F, 15.3% reduction), two androgen-dependent measures that rely heavily on steroid hormones produced by the testis [73, 74].

Finally, given the reduction in androgen-dependent measures in *Nr5a1-* Cre*;Nedd4^Flox/Flox^* testes, and the critical role of adult Leydig cells in steroid hormone synthesis, we next assessed the expression of genes whose products contribute to the regulation of steroidogenesis. These included hormones (*Insl3*), hormone receptors (*Ar* and *Lhr*) and gene products involved in hormone biosynthesis (*Star, Cyp11a1* and *Cyp17a1*). At P60, *Nr5a1-*Cre*;Nedd4^Flox/Flox^* testes had a significant reduction in expression of *Lhr* and *Cyp17a1,* while expression of *Ar, Star* and *Cyp11a1* remained unchanged. Surprisingly, however, mutant testes expressed *Insl3* transcripts at levels nearly twice that of littermate controls (Figure 5G). This provided compelling evidence that, beyond its role in Sertoli cell proliferation, NEDD4 contributes to the differentiation and function of the adult Leydig cell population within the postnatal testis.

## Discussion

Here we report previously uncharacterized roles of the ubiquitin ligase NEDD4 in the developing testis. While *Nr5a1*-Cre-driven ablation of *Nedd4* did not replicate the gonadal sex-reversal observed upon constitutive loss of this enzyme, its deletion in testicular somatic cells resulted in a significant reduction in postnatal testis weight. This phenotype stemmed from impaired Sertoli cell proliferation and defective adult Leydig cell differentiation.

### Inefficient Ablation of Nedd4 in Nr5a1-Cre;Nedd4^Flox/Flox^ Testes

Despite targeting *Nedd4* deletion in *Nr5a1*-positive cells, which are evident in the developing genital ridge as early as 10.2 dpc [60], *Nr5a1*-Cre;*Nedd4^Flox/Flox^* mice did not phenocopy the gonadal sex reversal observed in *Nedd4^-/-^* mice [48]. Investigation into NEDD4 ablation efficiency revealed that, while its expression was largely absent in most of somatic cells within the developing gonad, it persisted in the coelomic epithelium until at least 14.5 dpc (Figure S1C). This is consistent with previous reports showing incomplete deletion of genes such as *Fgfr2* [75], *Gata4* and *Fog2* [76] and *Gata4* and *Gata6* [77] when utilizing the same *Nr5a1*-Cre mouse line [52]. Despite continued expression of the deleted gene product within the coelomic epithelium, sex reversal phenotypes have been reported for *Fgfr2*, *Gata4* and *Sox9* [75, 76, 78]. These sex reversals, however, are often incomplete and are characterized by an inability to maintain the Sertoli cell fate. In contrast, earlier ablation of these genes with alternative Cre-drivers, causes complete sex reversal, often as a result of proliferation defects within the genital ridge, presumably resulting in an insufficient number of Sertoli cells to maintain the testicular fate [75, 76].

The complete sex reversal in XY *Nedd4^-/-^* gonads, but the absence of such a phenotype in *Nr5a1*-Cre;*Nedd4^Flox/Flox^*mice, supports the hypothesis that NEDD4 is essential for the proliferation of the coelomic epithelium and the establishment of the Sertoli cell stock, rather than for maintaining the male fate. This aligns with its oncogenic potential and its function in promoting Sertoli cell proliferation (Figure 3).

### Adrenal Cortex Dysgenesis in Nr5a1-Cre;Nedd4^Flox/Flox^ Mice

The adrenal dysgenesis observed in *Nr5a1-*Cre*;Nedd4^Flox/Flox^*mice is not unexpected, given the shared developmental origins of the adrenal cortex and the genital ridges, the male-to-female sex reversal observed in XY *Nedd4^-/-^* mice, and the similar adrenal and gonadal phenotypes reported in *Nr5a1* [79]*, Wt1* [80], *Cbx2* [81], *Cited2* [82], *Pbx1* [83], *Odd1* [84]*, Igf1r/Insr* [85] and *Six1/4* [86, 87] mutant mice. The same Cre driver has successfully generated adrenal gland phenotypes with other floxed alleles [64, 88–90].

Although *Nedd4* deletion in the adrenal cortex of *Nr5a1-*Cre*;Nedd4^Flox/Flox^*mice may be incomplete, as seen in the genital ridges, complete ablation of *Nedd4* may not be required to induce adrenal cortex dysgenesis.The adrenal cortex is highly sensitive to gene dosage, particularly for *Nr5a1*. Complete loss of this gene results in adrenal agenesis [79, 91–93], while heterozygous mice develop adrenal glands 12-fold smaller than wildtype littermates [94]. Indeed, given the near 50% reduction in *Nr5a1* transcripts at 11.5 dpc in XY *Nedd4^-/-^* gonads [48], it is likely that *Nr5a1* levels were sufficiently reduced in *Nr5a1-*Cre*;Nedd4^Flox/Flox^* adrenal cortexes to cause adrenal dysgenesis, such is the case in *Cbx2-* and *Cited2-*deficient mice [81, 82].

Interestingly, adrenocortical cells proliferation and steroidogenic activity increase later in development upon *Nr5a1* haploinsufficiency, perhaps as a means of compensating for earlier developmental defects [94–96], ultimately allowing fully functional adrenal gland in adulthood [97]. This could also be the case in *Nr5a1-* Cre*;Nedd4^Flox/Flox^* mice, given the progressive increase in the adrenocortical cell population between 14.5 and 18.5 dpc (Figure S2B). This may explain why these mice survive into adulthood, whereas more severe adrenocortical deficits in other models of adrenal dysgenesis results in early postnatal lethality [64, 89].

### NEDD4 Promotes Sertoli cell Proliferation

The absence of sex reversal in *Nr5a1-*Cre*;Nedd4^Flox/Flox^* mice provided a unique opportunity to explore the role of NEDD4 in the somatic cells of the postnatal testis, a function that remains largely uncharacterized due to the perinatal lethality [42] and gonadal sex reversal observed in XY *Nedd4^-/-^* mice.

The establishment of an adequate number of Sertoli cells in the mammalian testis is largely dependent on the length of the proliferative phase and the rate of division of these cells during development. In *Nr5a1-*Cre*;Nedd4^Flox/Flox^*testes, Sertoli cells exhibited reduced proliferative potential as early as 15.5 dpc (Figure 3), likely contributing to the reduction in postnatal testis-weight. Testis weight in *Nr5a1-* Cre*;Nedd4^Flox/Flox^* were reduced by 48.6% and 47.8% by P14 and P60, respectively (Figure 3B, G). However, in *Amh-*Cre*;Nedd4^Flox/Flox^*, testis weight was unchanged at P15 and only 29.7% reduced by P60 (Figure 4B). This likely stems from the timing of *Nedd4* ablation. The *Nr5a1-Cre* line deletes genes as early as 11.5 dpc, whereas *Amh-*Cre ablation occurs around 15 dpc [52, 56]. Consequently, Sertoli cell proliferation of *Nr5a1-*Cre*;Nedd4^Flox/Flox^* Sertoli cells experience earlier disruption in proliferation than those of *Amh-*Cre*;Nedd4^Flox/Flox^*Sertoli cells, reducing the total rounds of division, and leading to a more pronounced reduction of Sertoli cell number and testis weight. Interestingly, *Nr5a1-*Cre*;Nedd4^Flox/Flox^*testes were already smaller by 12.5 dpc (Figure 1), suggesting that this reduction in proliferative potential arose early in testis differentiation, though not early enough to cause gonadal sex reversal. Furthermore, the additional defects in adult Leydig cell differentiation (Figure 5) may further exacerbate the reduced testis size in *Nr5a1-*Cre*;Nedd4^Flox/Flox^* mice compared to *Amh-*Cre*;Nedd4^Flox/Flox^*mice.

The role of NEDD4 in Sertoli cell proliferation aligns with its previously reported contributions to genital ridge proliferation [48], and its purported role as an oncogene [67–69, 98–100]. Our findings indicate that NEDD4 modulates the PI3K-AKT signaling pathway, which was diminished in *Nedd4-*mutant testes (Figure 3J, K). Given that numerous signaling cascades converge on AKT activation, many of which are critical for Sertoli cell proliferation and homeostasis [101–103], NEDD4 likely facilitates testis growth via this pathway. Diminished AKT activation in *Nr5a1-*Cre*;Nedd4^Flox/Flox^*testes, however, occurs in the absence of ectopic PTEN, a NEDD4 substrate known to drive similar phenotypes upon loss of *Nedd4*. These observations add to the complexity of the NEDD4 and PTEN relationship. While some *in vivo* studies supports negative regulation of PTEN by NEDD4, with PTEN levels increasing upon *Nedd4* deletion [104–106] and decreasing with ectopic NEDD4 expression [106], PTEN stability and localization remain largely unaffected in various *Nedd4*-deficient tissues, including fibroblasts [42, 107], T cells [43, 108], neurons [109] and spermatogonial stem cells [49]. Our study (Figure 3J), similarly found no change in PTEN expression, suggesting that the regulation of PTEN by NEDD4 may be context dependent or that redundant mechanisms maintain PTEN stability and localization in the absence of NEDD4. Indeed, this may be true of many *in vitro* identified NEDD4 substrates as known substrates Beclin1 [110], p62 [111], LC3 [112, 113] and PDCD6IP [50, 114, 115] remain unchanged in *Nedd4-*mutant testes (Figure S3).

Beyond its role in PTEN regulation, NEDD4 is required for animal growth and this has been linked to the control of IGF-1 and insulin signaling via the PI3K-AKT pathway. Ubiquitination of GRB10 and IRS2 by NEDD4 have been implicated in such regulation [42, 116]. In keeping with this, insulin and IGF-1 signaling is diminished upon loss of *Nedd4.* Phenotypes resembling those of *Nedd4-*mutants arise upon dual ablation of the insulin receptor (*Insr*) and insulin-like growth factor 1 receptor (*Igf1r*). In this way XY *Insr^-/-^;Igf1r^-/-^* mice exhibit diminished proliferation of somatic progenitors within the genital ridges, XY gonadal sex reversal and perturbed adrenal development [85]. *Insr;Igf1r*-mutants also exhibit significant reductions in testis weight, owing to reduced Sertoli cell proliferation [40], though this is rescued upon ablation of *Pten* [70]. Given the known regulation of insulin and IGF-1 signaling by NEDD4, and the importance of this signaling axis across multiple levels of testis differentiation, perturbed insulin and IGF-1 signaling may underlie the phenotypes observed in *Nedd4* mutants, particularly given the marked reduction in pAKT observed in mutant testes (Figure 3). While the precise mechanism of NEDD4’s control over the PI3K-AKT axis in murine Sertoli cells remains to be elucidated, NEDD4 is known to interact with multiple components along this signaling cascade [117–120]. Investigating how NEDD4 modulates these interactions should prove illuminating to further our understanding of Sertoli cell dynamics.

### NEDD4 Regulates Adult Leydig Cell Differentiation and Steroidogenesis

Beyond its roles in adrenal cortex development and Sertoli cell proliferation, NEDD4 is essential for proper adult Leydig cell differentiation. In *Nr5a1-*Cre*;Nedd4^Flox/Flox^* testes, the transition from progenitor to immature adult Leydig cells was notably disrupted (Figure 5C). This defect contributed to reduced androgen-sensitive parameters, such as anogenital distance and seminal vesicle weight (Figure 5D, E, F), alongside diminished steroidogenesis (Figure 5G).

Interestingly, while other models of adult Leydig cell dysfunction show a near absence of *Cyp11a1, Cyp17a1, Star, Insl3* and *Lhr* [121], of these, only *Cyp17a1* and *Lhr* were significantly decreased in *Nr5a1-*Cre*;Nedd4^Flox/Flox^* testes, suggesting a more nuanced role for NEDD4 in steroidogenesis. Notably, *Star* and *Cyp11a1* act early in steroidogenesis. STAR regulates cholesterol transport into mitochondria while CYP11A1 converts cholesterol to pregnenolone [122]. CYP17A1 acts downstream of these proteins, ultimately aiding in the conversion of pregnenolone and progesterone to their major products: estradiol, testosterone, DHEA and cortisol [123]. Given the observed expression data, it is likely that NEDD4, if it were intrinsically involved in this process, would exert its influence downstream of STAR and CYP11A1 and upstream of CYP17A1. *Hsd17b3* transcripts are also diminished in *Nedd4-*mutant testes which, itself, acts downstream of CYP17A1, further supporting this hypothesis.

### Disrupted LHR Signaling Contributes to Impaired Adult Leydig Cell Differentiation

The downregulation of *Lhr* in *Nr5a1-*Cre*;Nedd4^Flox/Flox^*testes provides further insight into the Leydig cell phenotype observed upon *Nedd4* deletion. LHR signaling is essential for adult Leydig cell differentiation, yet dispensable for the development of fetal Leydig cells. While *Lhr* is expressed in fetal Leydig cells [124], *Lhr*-deficient mice exhibit normal Leydig cell development at fetal and neonatal stages [125]. In contrast, adult *Lhr*^-/-^ mice exhibit significant reductions in Leydig cell numbers, anogenital distance, testis weight and seminal vesicle weight [125, 126]. Surviving Leydig cells in *Lhr*-deficient testes are also functionally immature, as evidenced by a decrease in cytoplasmic volume [127]. These observations align with the unaffected differentiation of fetal Leydig cells in *Nr5a1-*Cre*;Nedd4^Flox/Flox^* testes (Figure 2) and the postnatal Leydig cell dysfunction observed later (Figure 5). This suggests that impaired LHR signaling contributes to the Leydig cell defects observed upon ablation of *Nedd4*. However, since *Lhr* levels are only reduced by approximately 28% in *Nr5a1-*Cre*;Nedd4^Flox/Flox^*testes, rather than absent in *Lhr*-deficient mice, the Leydig cell phenotype in *Nr5a1-*Cre*;Nedd4^Flox/Flox^* is comparatively milder. While a thorough characterization of *Lhr* heterozygous mice is lacking, in which *Lhr* transcript levels are significantly reduced, the serum hormone profile of these mice is altered and Leydig cells do appear morphologically different to wildtype littermates [125, 126], perhaps suggesting that a mere reduction in *Lhr* expression is sufficient to induce Leydig cell associated phenotypes, as is observed in this study. *Cyp17a1* expression has also been shown to be luteinizing hormone (LH) dependent, yet independent of testosterone, unlike other genes encoding steroidogenic components [128], suggesting that the reduction in *Cyp17a1* may merely be a consequence of reduced *Lhr* expression. This may further explain why *Cyp17a1* is reduced in *Nr5a1-* Cre*;Nedd4^Flox/Flox^*testes while *Cyp11a1* and *Star* remain unchanged (Figure 5G).

Curiously, despite the observed downregulation of *Cyp17a1* and *Lhr*, *Nr5a1-* Cre*;Nedd4^Flox/Flox^* testes showed a significant increase in *Insl3* expression. This finding is intriguing, as INSL3 is widely recognized as a biomarker of Leydig cell differentiation and function [129, 130]. It is perhaps paradoxical that a mouse model exhibiting perturbed Leydig cell function exhibit increased levels of *Insl3* transcripts. One possible explanation is that the reduction in *Lhr* expression triggers a compensatory increase in circulating LH. Indeed, gonadotropins (luteinizing hormone and human chorionic gonadotropin) are known to upregulate *Insl3* expression [131] and *Lhr* deficient mice have increased circulating LH [125, 126, 132]. In this way, it is feasible that the observed reduction in *Lhr* in *Nr5a1-*Cre*;Nedd4^Flox/Flox^* testes results in a compensatory increase in circulating LH which subsequently upregulates *Insl3*. How increased levels of *Insl3* affect Leydig cell function, however, remains to be seen. While transgenic *Insl2/Insl3* expression was capable of rescuing the cryptorchidism of *Insl3^-/-^* mice, the transient increase in *Insl3* expression in transgenic mice had no impact on male fertility [133], and no testicular abnormalities were reported.

In conclusion, this study highlights the distinct and multifaceted role of the ubiquitin ligase NEDD4 in murine testis development. We demonstrate that NEDD4 plays a crucial role in the early establishment of the adrenogonadal primordium, with its retention in the gonadal coelomic epithelium being sufficient to rescue the gonadal sex reversal observed in constitutive knockouts. However, this is not the case in the adrenal cortex, where *Nedd4* ablation results in adrenal dysgenesis, underscoring the differential sensitivity of these tissues to *Nedd4* ablation. Consistent with its reported role as an oncogene, we have shown that NEDD4 promotes Sertoli cell proliferation, through the modulation of the PI3K-AKT signaling pathway. NEDD4 also ensures proper differentiation of adult Leydig cells and may contribute to murine steroidogenesis. Overall, this study establishes that NEDD4 functions at multiple levels to orchestrate testis development, reinforcing the importance of ubiquitination, and post-translational modifications in the differentiation of the mammalian testis.

## Acknowledgments

The authors would like to thank Dr Hiroshi Kawabe (Max-Planck-Institute of Experimental Medicine) for kindly providing the *Nedd4^flox/flox^* mice, the staff of University of Melbourne’s Biomedical Sciences Animal facility for mouse colony maintenance and animal husbandry, the Melbourne Histology Platform of the University of Melbourne’s School of Biomedical Sciences for tissue processing and the University of Melbourne’s Biological Optical Microscopy Platform for use of their confocal microscopes.

## Grants or Fellowships

This work was supported by an Australian Research Council Discovery Grant (150101448) awarded to D.W. and S.K. and partly supported by a Monash University Biomedicine Discovery Institute Early Career Award awarded to S.W. S.K. was supported by a National Health and Medical Research Council of Australia Investigator Grant (GNT 2007739). S.N. and Y.N. received funding from the Swiss National Science Foundation under grants 31003A_173070 and 310030_200316.

---

**Supplementary Figure 1.**
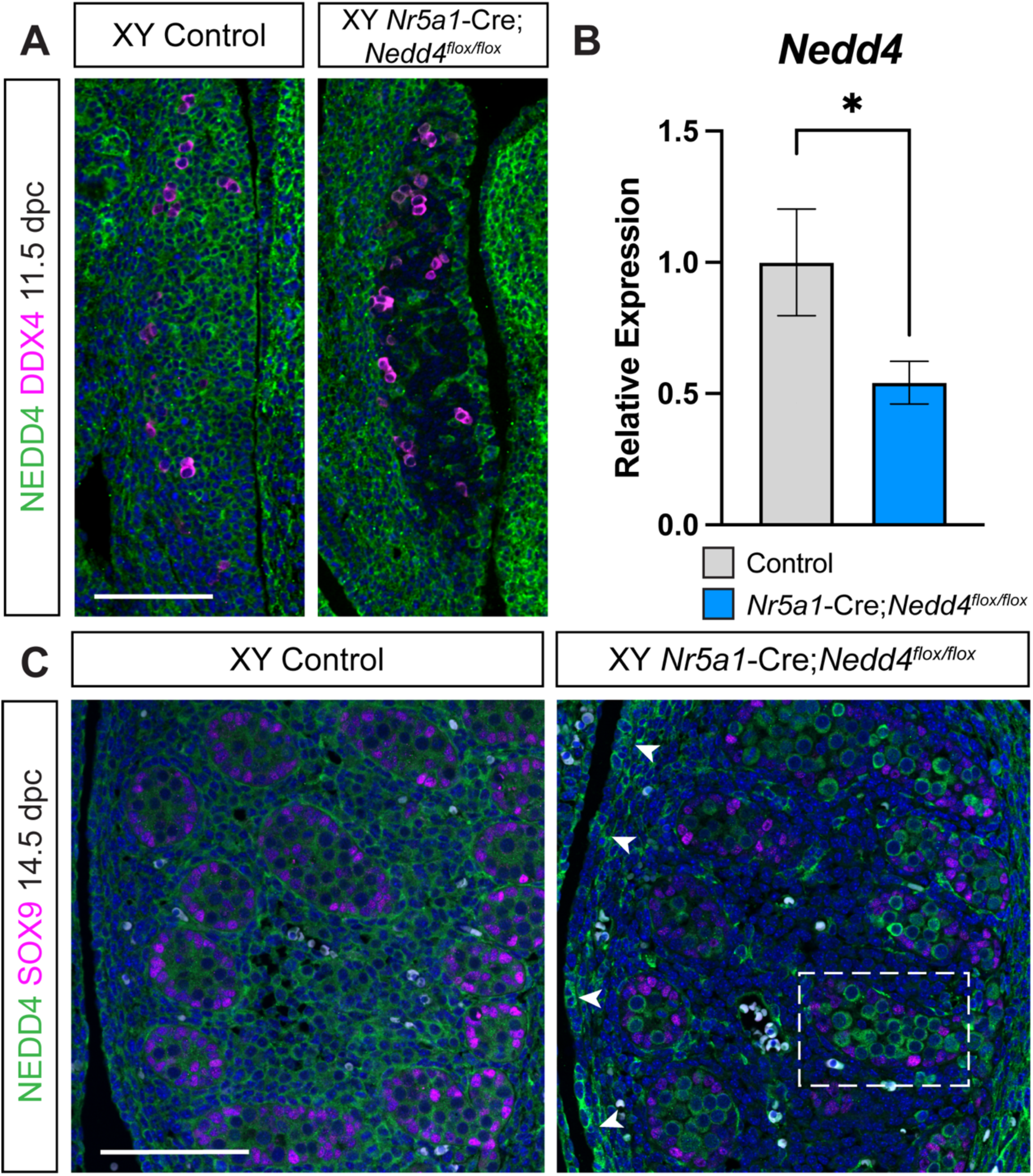
Ablation of NEDD4 in Somatic Cells of the Testis. **A)** Section immunofluorescence on 11.5 dpc XY *Nr5a1-*Cre*;Nedd4^Flox/Flox^*testes alongside Cre-negative XY controls stained for NEDD4 (green) and germ cell marker DDX4 (magenta). Scale bar = 100 μm. **B**) RT-qPCR analysis of *Nedd4* expression at 14.5 dpc on XY *Nr5a1-*Cre*;Nedd4^Flox/Flox^*gonads (blue, n=5) and XY Cre-negative littermate controls (grey, n=5). Values are normalized to *Sdha* and are expressed relative to controls. Mean ± SEM; t-test; * = p < 0.05. **C**) Section immunofluorescence on 14.5 dpc XY *Nr5a1-*Cre*;Nedd4^Flox/Flox^* testes alongside Cre-negative XY controls stained for NEDD4 (green) and Sertoli cell marker SOX9 (magenta). Scale bar = 100 μm.

**Supplementary Figure 2.**
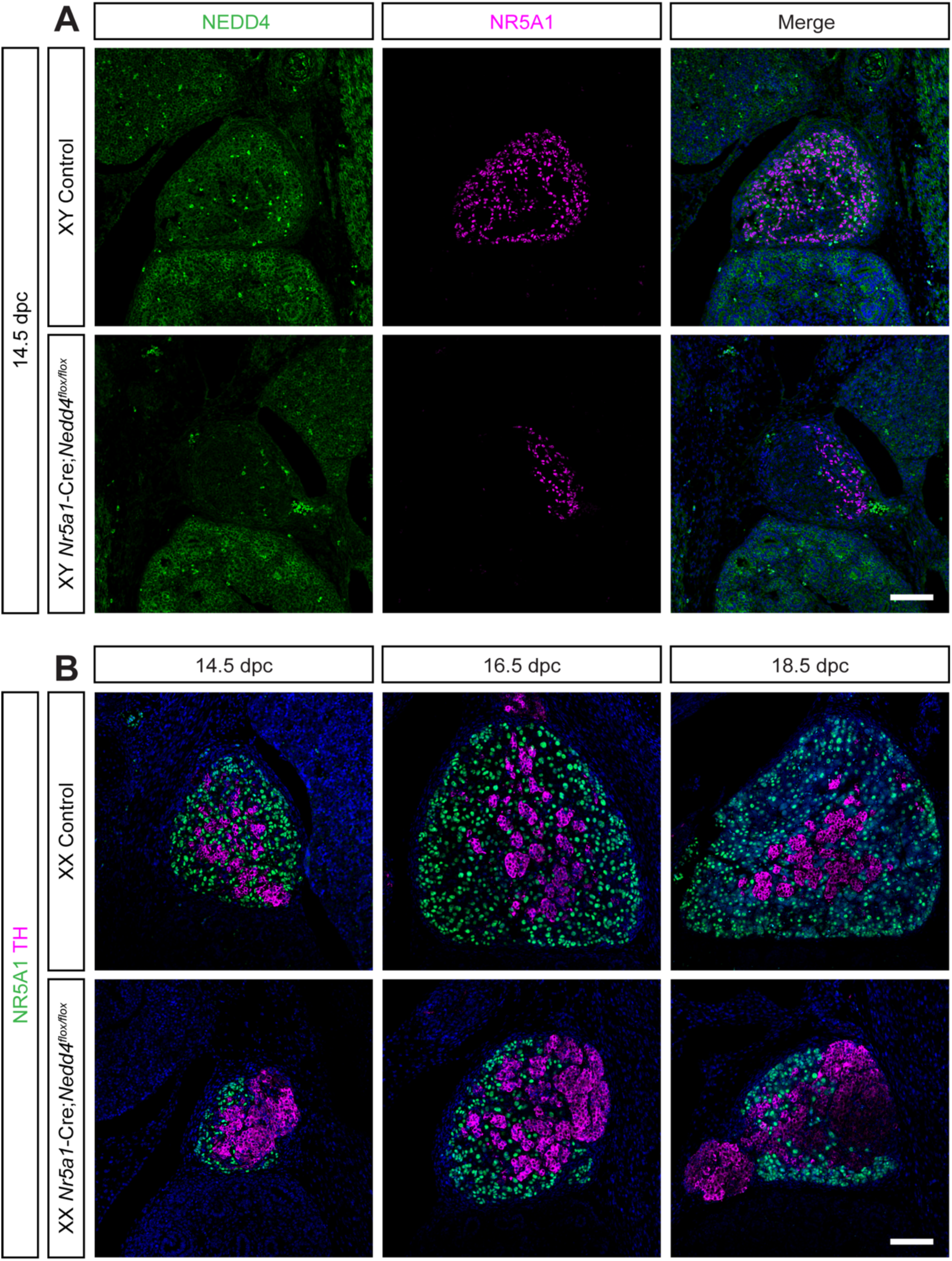
Adrenal Cortex Dysgenesis in *Nr5a1-Cre;Nedd4^flox/flox^* Mice. **A)** Section immunofluorescence on 14.5 dpc *Nr5a1-*Cre*;Nedd4^flox/flox^* adrenal glands alongside Cre-negative littermate controls stained for NEDD4 (green) and NR5A1 (magenta) merged with nuclear stain DAPI (blue). **B**) Section immunofluorescence on *Nr5a1-*Cre;*Nedd4^flox/flox^*and Cre-negative littermate controls at 14.5, 16.5 and 18.5 dpc stained for the steroidogenic, adrenal cortex marker NR5A1 (green) and the chromaffin cell, adrenal medulla marker TH (magenta) alongside nuclear stain DAPI (blue). Scale bars = 100 μm.

**Supplementary Figure 3.**
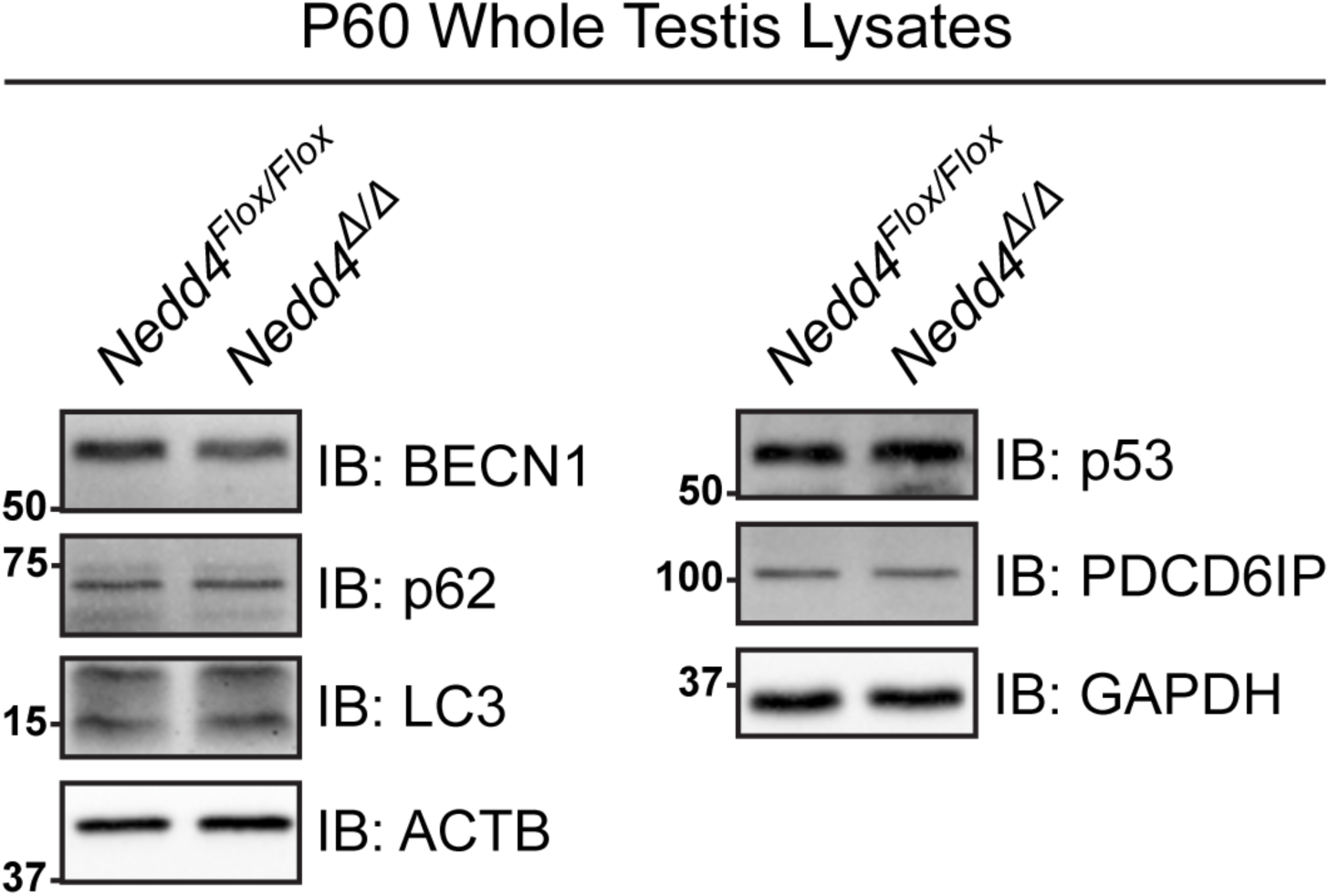
Immunoblots. **A**) Immunoblots of known NEDD4 substrates on P60 *Nr5a1-*Cre*;Nedd4^flox/flox^* testis lysates (*Nedd4^Δ/Δ^*, right) alongside Cre-negative littermate controls. Membranes were probed for Beclin 1 (BECN1), p62 and LC3 using Beta Actin (ACTB) as a loading control. Membranes were also probed p53 and PDCD6IP, with GAPDH serving as a loading control.

**Supplementary Table 1.**
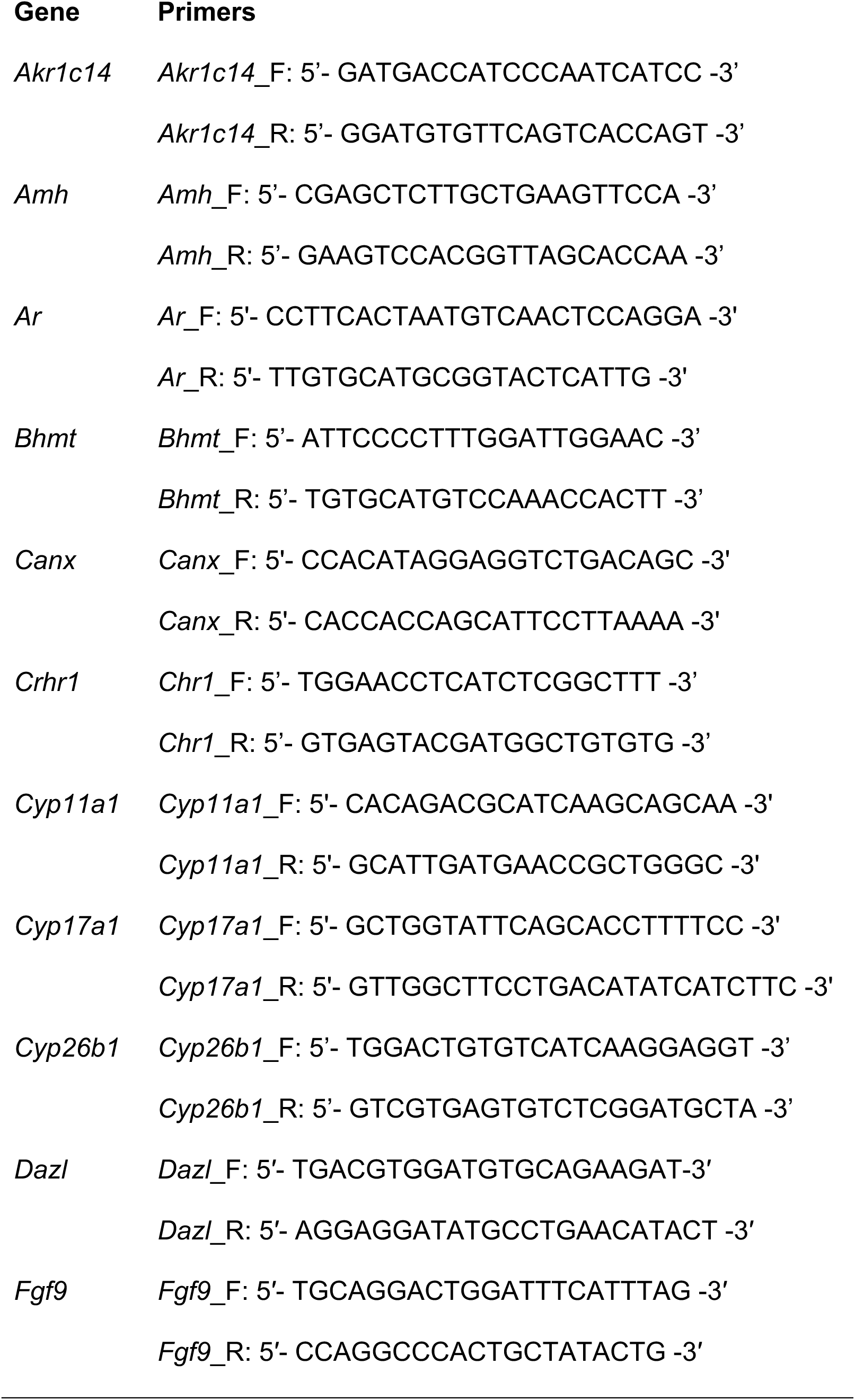
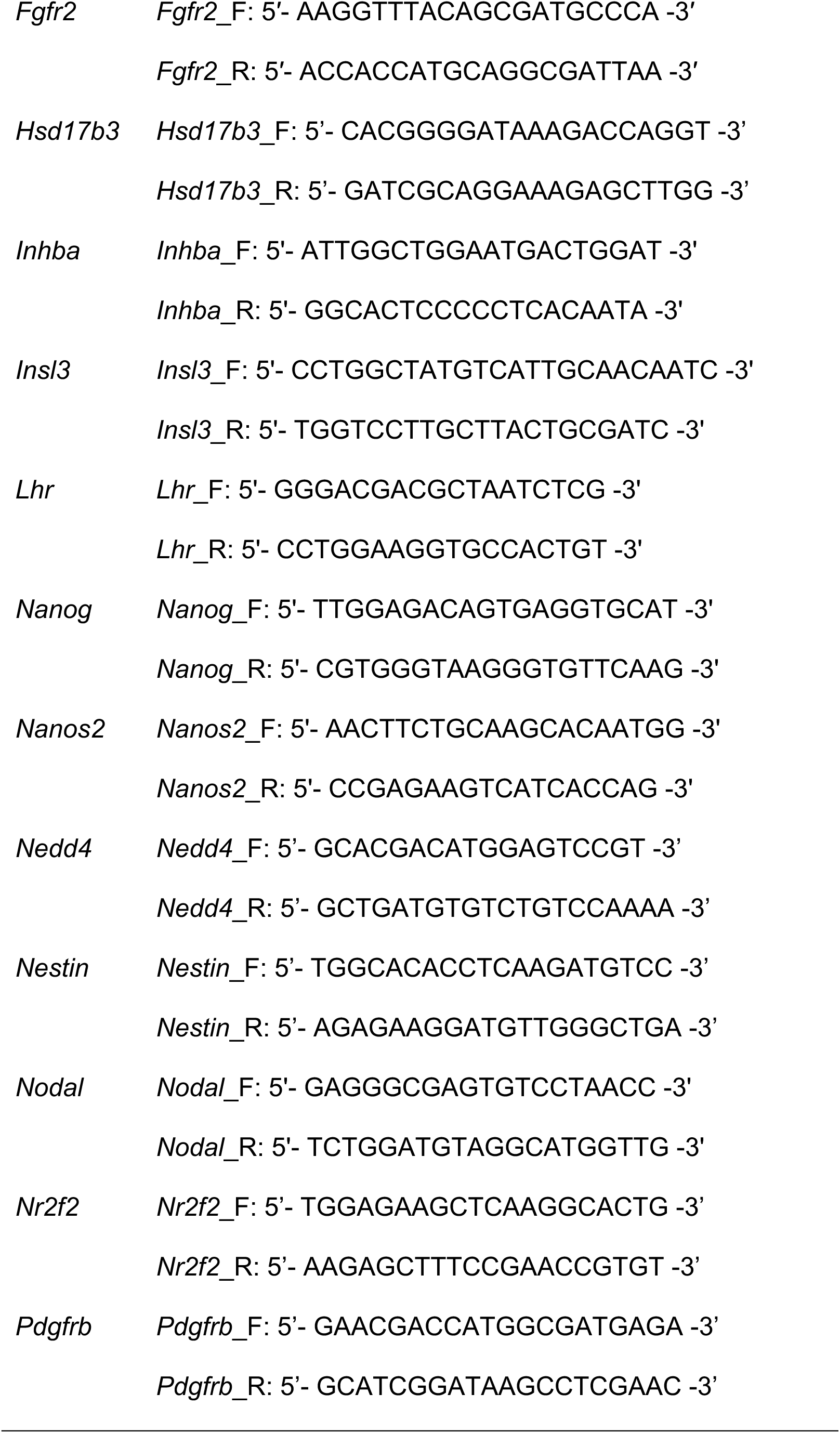
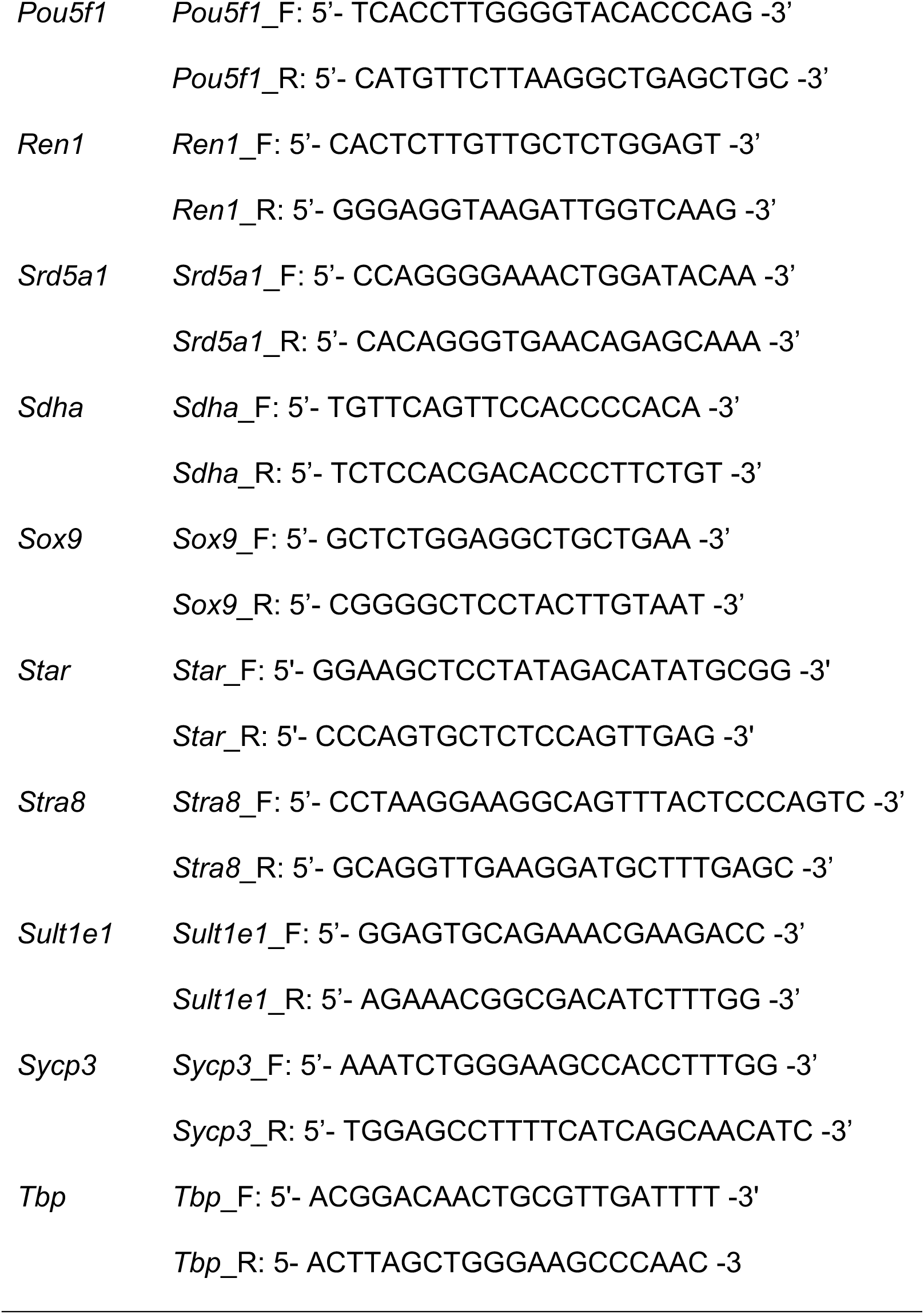
RT-qPCR Primers Gene Primers.

